# Association of NOTCH3 with Elastic Fiber Dispersion in the Infrarenal Abdominal Aorta of Cynomolgus Monkeys

**DOI:** 10.1101/2023.03.04.530901

**Authors:** Sohei Ito, Naofumi Amioka, Michael K. Franklin, Pengjun Wang, Ching-Ling Liang, Yuriko Katsumata, Lei Cai, Ryan E. Temel, Alan Daugherty, Hong S. Lu, Hisashi Sawada

**Affiliations:** Saha Cardiovascular Research Center, College of Medicine; Saha Aortic Center, College of Medicine, University of Kentucky, KY; Department of Physiology, College of Medicine, University of Kentucky, KY; Department of Biostatistics, College of Public Health, University of Kentucky, KY; Sanders-Brown Center on Aging, University of Kentucky, KY

## Abstract

**Background:** The regional heterogeneity of vascular components and transcriptomes is an important determinant of aortic biology. This notion has been explored in multiple mouse studies. In the present study, we examined the regional heterogeneity of aortas in non-human primates.

**Methods:** Aortic samples were harvested from the ascending, descending, suprarenal, and infrarenal regions of young control monkeys and adult monkeys provided with high fructose for 3 years. The regional heterogeneity of aortic structure and transcriptomes was examined by histological and bulk RNA sequencing analyses.

**Results:** Immunostaining of CD31 and αSMA revealed that endothelial and smooth muscle cells were distributed homogeneously across the aortic regions. In contrast, elastic fibers were less abundant and dispersed in the infrarenal aorta compared to other regions and associated with collagen deposition. Bulk RNA sequencing identified a distinct transcriptome related to the Notch signaling pathway in the infrarenal aorta with significantly increased *NOTCH3* mRNA compared to other regions. Immunostaining revealed that NOTCH3 protein was increased in the media of the infrarenal aorta. The abundance of medial NOTCH3 was positively correlated with the dispersion of elastic fibers. Adult cynomolgus monkeys provided with high fructose displayed vascular wall remodeling, such as smooth muscle cell loss and elastic fiber disruption, predominantly in the infrarenal region. The correlation between NOTCH3 and elastic fiber dispersion was enhanced in these monkeys.

**Conclusions:** Aortas of young cynomolgus monkeys display regional heterogeneity of their transcriptome and the structure of elastin and collagens. Elastic fibers in the infrarenal aorta are dispersed along with upregulation of medial NOTCH3.

**HIGHLIGHTS:**

- The present study determined the regional heterogeneity of aortas from cynomolgus monkeys.
- Aortas of young cynomolgus monkeys displayed region-specific aortic structure and transcriptomes.
- Elastic fibers were dispersed in the infrarenal aorta along with increased NOTCH3 abundance in the media.

**GRAPHIC ABSTRACT:** 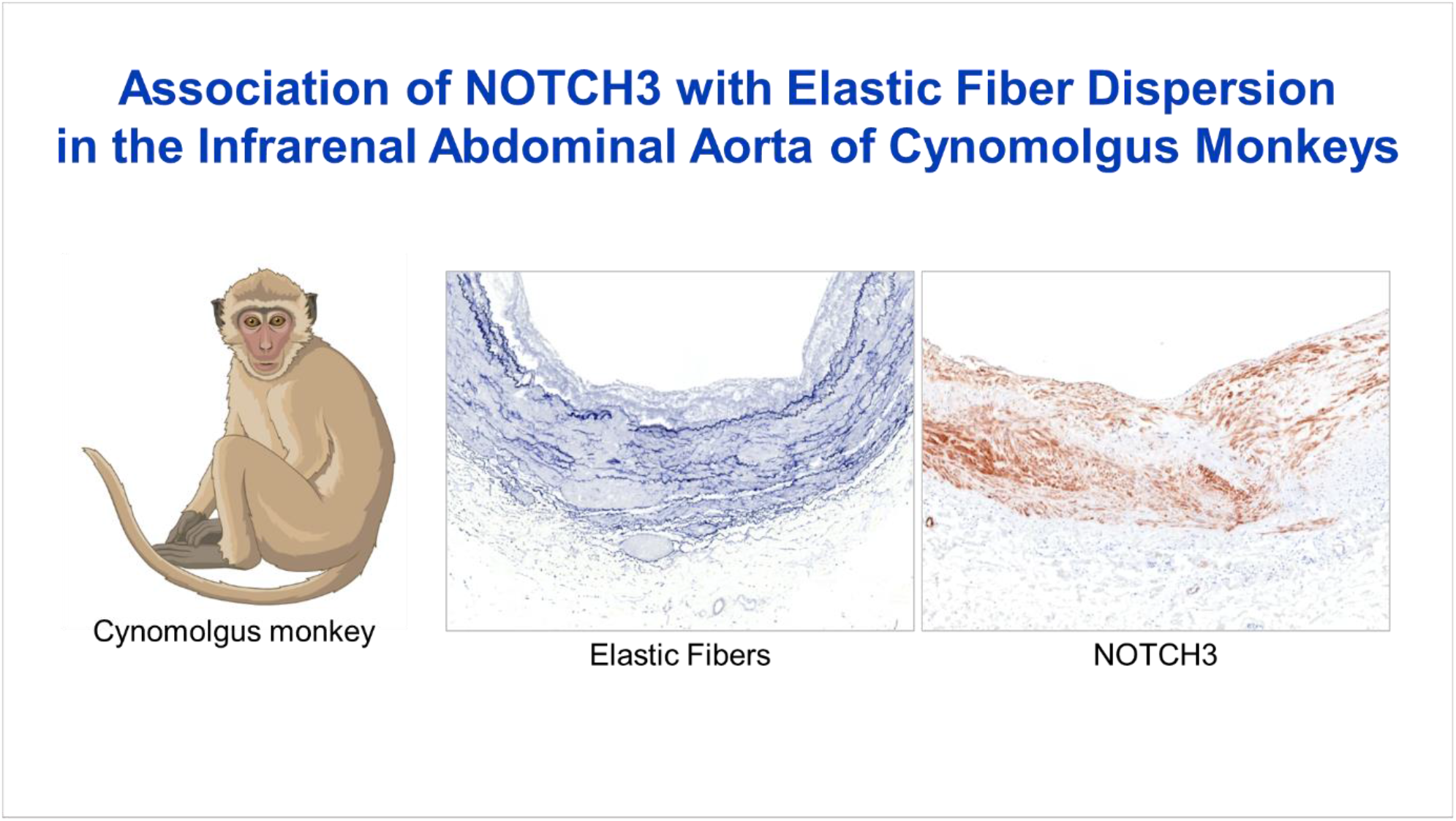

## INTRODUCTION

The aorta is the largest vessel in the body that supplies oxygenated blood to the systemic circulation. The aortic wall is composed of three tunicae: intima, media, and adventitia, which primarily contain endothelial cells (ECs), smooth muscle cells (SMCs), and fibroblasts, respectively. These resident cells play pivotal roles in maintaining the integrity of the aortic wall. In addition to these cells, the extracellular matrix (ECM) is also an important structure in the aortic wall. Aortic cells and ECM form a laminar structure cylindrically that confers structural strength and elasticity to the vascular wall.^1, 2^ Thus, it is vital to understand the precise roles of these components for advancing our knowledge of aortic biology.

The aorta is commonly described in four major regions: ascending and descending thoracic regions and suprarenal and infrarenal abdominal regions. Although the laminar structure of the vascular wall is superficially homogeneous throughout the aorta, there is evolving evidence that aortic functions are heterogeneous across the regions. For instance, an in vitro study using chick cells demonstrated that SMCs in the aortic arch are more proliferative than those in the infrarenal abdominal aorta.^3^ In mice, angiotensin II infusion induced cellular hyperplasia in the ascending aorta but cellular hypertrophy in other regions.^4^ Of note, only the infrarenal abdominal region of mouse aorta constricts in response to angiotensin II.^5^ These studies support the notion that different aortic regions display distinct biological behaviors. Despite multiple studies in animal models, this hypothesis has not been fully examined in humans. A recent study using aortic images by computational tomography revealed higher vascular stiffness in the infrarenal abdominal aorta than in the ascending aorta.^6^ Another study demonstrated the regional variance of collagen and elastin contents histologically using human aortic samples from aged (67-92 years old) subjects.^7^ To further understand the regional heterogeneity, it is important to determine histological differences and their molecular basis in young subjects without pathological changes. However, the limited availability of freshly harvested human samples, especially aortas from healthy young individuals, is a significant impediment to the determination of the regional specificity of aortic biology under physiological conditions.

Mice have been used as a common species in preclinical studies to investigate vascular physiology, including the regional heterogeneity of the aorta.^4, 5^ However, mice may not mimic the human vascular system. Aortic size and hemodynamics are considerably different between mice and humans.^6^ Although blood pressure is similar between the species, heart rate is 10 times higher (human: 60-80, monkey: 100-170, mouse: 500-700 beats per minute) in mice and their elastic laminar layers in the aorta are significantly less than in humans (3-5 layers vs >20 layers in the infrarenal aortic region).^8-13^ Given these differences, studies using mouse models often require validation using human samples for translational insights. In contrast, non-human primates have similarities with humans in these parameters. Although non-human primates have modestly higher heart rate than humans, their aortic structure, including the number of elastic laminar layers, are comparable to humans. Furthermore, there is a genetic, metabolic, and physiologic proximity between humans and non-human primates.^14^ Therefore, studies using non-human primates should provide more translational insights into understanding aortic physiology.

In the present study, bulk RNA sequencing analyses and histological were performed in aortas of young healthy cynomolgus monkeys to understand the transcriptomic heterogeneities across the aortic regions under physiological conditions. The regional heterogeneity of aortic histology was also determined in adult monkeys with atherosclerotic lesions.

## MATERIAL AND METHODS

### Non-human Primates

Male and female Mauritian cynomolgus monkeys were purchased from DSP Research Services (3 and 5 years old, n=3-4). Monkeys were initially fed a standard nonhuman primate laboratory diet ad libitum (Teklad 2050, Envigo) and given access to water ad libitum. Male monkeys were housed singly from 8:00 - 15:00 each day and received a high fructose diet (**Supplemental Table 1**) in the morning and afternoon. In place of water, a high fructose drink (**Supplemental Table 1**) was provided ad libitum to the male monkeys. High fructose diet and drink were fed for 36 months. A standard laboratory diet was fed throughout the study to the female monkeys. The room temperature and humidity were maintained at 68 to 74°F and 50 to 60%, respectively. Monkeys were maintained on a 14:10 hour light:dark cycle. Male and female monkeys were euthanized at 8 and 3 years old, respectively. The monkeys were fasted overnight and sedated with ketamine (10 mg/kg, im) and isoflurane (3-5% vol/vol induction, 1-2% vol/vol maintenance). After an adequate depth of anesthesia was established by lack of physical response, the inferior vena cava was exposed and cut for exsanguination. A 16-gauge needle was inserted into the left ventricle of the heart and saline was perfused to flush the body of blood. The euthanasia method was AVMA acceptable. Aortic tissues were harvested from each region (**Supplemental Figure 1**): the middle ascending aorta (Asc), the middle descending thoracic aorta (Dsc), the suprarenal abdominal aorta proximal to the celiac artery (Sup), and the distal infrarenal abdominal aorta proximal to the bifurcation of iliac arteries (Inf). Aortic samples were either snap-frozen in liquid nitrogen for bulk RNA sequencing or immersed in paraformaldehyde (4% wt/vol, P6148, Sigma-Aldrich) for histological analyses. All protocols were approved by the University of Kentucky IACUC (#2020-3596) in accordance with the National Institutes of Health guidelines.

### Histological Staining

Fixed aortas were placed in ethanol (70% vol/vol) for 24 hours, embedded into paraffin, and cut into 5 µm sections. Subsequently, paraffin-embedded sections were deparaffinized using limonene (183164, Sigma-Aldrich) followed by 2 washes with ethanol (100%) and 1 wash with Automation Buffer (GTX30931, GeneTex).

For immunostaining, deparaffinized sections were incubated with H_2_O_2_ (1%; H325-500, Fisher Scientific) for 2 minutes at 40°C. Antigen retrieval (HK547-XAK, BioGenex, 20 minutes at 98°C) was applied for CD31, NOTCH3, Ki67, and CD68 staining. Non-specific binding of primary antibodies was blocked by goat serum (2.5% vol/vol; MP-7451, Vector laboratories) for 20 minutes at room temperature. Sections were next incubated with rabbit alpha-smooth muscle actin (α-SMA) antibody (2 µg/mL, ab5694, abcam) for 30 minutes at room temperature, rabbit CD31 antibody (3 µg/mL, ab28364, abcam), rabbit COL1A1 antibody (3 μg/mL, ab138492, abcam), rabbit Ki67 antibody (0.03 µg/mL, ab16667, abcam), mouse anti human CD68 antibody (0.5 µg/mL, 916104, BioLegend), or rabbit NOTCH3 antibody (1 µg, ab23426, abcam) for overnight at 4°C. Sections were also incubated with a non-immune rabbit IgG antibody (I8140, Sigma-Aldrich) as negative controls. Goat anti-rabbit IgG conjugated with horseradish peroxidase (30 minutes, MP-7451, Vector laboratories) was used as a secondary antibody. ImmPACT® NovaRed (SK4805, Vector) was used as a chromogen. Hematoxylin (26043-05, Electron Microscopy Sciences) was used for counterstaining. Slides were coverslipped with Permanent Mounting Medium (H-5000, Vector).

Movat’s pentachrome staining was performed using Movat’s Pentachrome method for Connective Tissue kits (k042, Poly Scientific R×D) according to the manufacturer’s protocol. Verhoeff’s iron hematoxylin was performed as described previously.^15, 16^ Briefly, deparaffinized sections were incubated with Verhoeff’s iron hematoxylin (iodine: 5.3 g/L, potassium iodide: 10.7 g/L, hematoxylin: 2.8% wt/vol, ferric chloride: 2.2% wt/vol) for 20 minutes. Subsequently, the sections were differentiated with ferric chloride (2% wt/vol) for 2 minutes and washed with distilled water. After dehydration, the sections were coverslipped with Permount (SP100, Fisher Scientific). For picrosirius red (PSR) staining, deparaffinized sections were incubated with phosphomolybdic acid (0.2% vol/vol; HT153, Sigma-Aldrich) for 5 minutes followed by staining with picrosirius red and fast green solution: Direct Red 80 (0.1% vol/vol; 365548, Sigma-Aldrich), Fast Green FCF (0.1% wt/vol; F7258, Sigma-Aldrich) in saturated picric acid (1.3% wt/vol; P6744, Sigma-Aldrich), for 60 minutes at room temperature. The slides were then rinsed with acidified water (0.1 N hydrochloride acid, H2505, Aqua Solutions Inc), dehydrated through increasing concentration of ethanol, incubated with three changes of xylene, and coverslipped with DEPEX (50-247-470, Thermo Fisher Scientific).

### Histological Measurements

Histological images were captured using either Eclipse Ni with DS-Ri2 (Nikon) or AxioScan 7 (Zeiss). The images were analyzed using either NIS-Elements AR software (ver 5.11.03, Nikon) or Zen Blue edition (ver 3.3, Zeiss). Internal perimeters were measured by tracing the internal elastic lamina in Verhoeff iron hematoxylin stained sections. Medial thickness was defined as the distance from the inner to external elastic laminae. Medial area was defined as the extent between the internal and external elastic laminae and measured using low magnification pictures that captured the entire aortas. Elastic lamellar thickness and interlamellar distance were also measured using Verhoeff iron hematoxylin staining images. Collagen and NOTCH3 positive areas in the media of picrosirius red and immunostained sections were measured using the following color threshold: collagen: R: 0-255, G: 0-255, B: 0-160; NOTCH3: R: 0-255, G: 0-255, B: 0140. These measurements were performed in four orthogonal locations of each section with the most thickened location as a reference point (**Supplemental Figure 2**) and means of the four locations were calculated. All measurements were verified by a member of the laboratory who was blinded to the identification of study groups.

### Western Blot Analyses

Aortic tissues were homogenized in Cell Lysis buffer (9803, Cell Signaling Technology) with a protease inhibitor cocktail (P8340, Sigma-Aldrich) using Kimble Kontes disposable Pellet Pestles (Z359971, DWK Life Science LLC.,). Protein lysate from HT-1080 cells incubated with and without hTGF-β3 for 30 minutes was used as positive and negative controls, respectively (12052, Cell Signaling Technology). Protein concentrations were determined using a DC assay kit (5000111, Bio-Rad). Equal masses of protein per sample (20 µg) were resolved by SDS-PAGE (10% wt/vol) and transferred electrophoretically to PVDF membranes. After blocking, antibodies against the following proteins were used to probe membranes: Collagen type I (0.9 µg/mL, ab138492, abcam), p-SMAD2 (0.1 µg/mL, 3108S, Cell Signaling Technology), SMAD2 (0.1 µg/mL, 3122S, Cell Signaling Technology), p-ERK (0.1 µg/mL, 9101S, Cell Signaling Technology), ERK (0.1 µg/mL, 9102S, Cell Signaling Technology), and β-actin (0.3 µg/mL, A5441, Sigma-Aldrich). Membranes were incubated with either goat anti-rabbit (1.0 µg/mL, PI-1000, Vector Laboratories) or goat anti-mouse secondary antibodies (0.3 µg/mL, A2554, Sigma-Aldrich). Immune complexes were visualized by chemiluminescence (34080, Thermo Fisher Scientific) using a ChemiDoc (12003154, BioRad) and quantified using Image Lab software (v6.0.0, BioRad).

### Gelatin Zymography

Aortic samples were homogenized in Cell Lysis buffer and aortic protein (10 µg) was dissolved in zymogram sample buffer (786-483, G-Biosciences). Subsequently, samples were loaded onto zymography gels (ZY00102BOX, Thermo Fisher Scientific) and run with SDS running buffer for 90 minutes (LC2675, Thermo Fisher Scientific). Gels were renatured in Zymogram Renaturing Buffer (LC2670, Thermo Fisher Scientific) for 30 minutes at room temperature and incubated with Zymogram Developing Buffer (LC2671, Thermo Fisher Scientific) for 30 minutes at 37°C. Gels were incubated further with fresh developing buffer for 36 hours at 37°C. Then, gels were washed with dH_2_O and stained with SimplyBlue SafeStain (LC6060, Thermo Fisher Scientific) for 20 minutes. A ChemiDoc system was used to acquire images of the stained gels.

### Bulk RNA sequencing

Aortic samples were incubated with RNAlater solution (#AM7020, Thermo Fisher Scientific) for 12 hours. Subsequently, RNA was extracted using RNeasy Fibrous Tissue Mini kits (#74704, Qiagen). mRNA samples were shipped to Novogene for bulk mRNA sequencing. cDNA library was generated from total mRNA (1 µg) using NEBNext UltraTM RNA Library Prep Kits for Illumina (New England BioLabs). cDNA libraries were sequenced by NovaSeq 6000 (Illumina) in a paired-end fashion to reach more than 1.5M reads. The Qscore of over 50% bases of the read is ≤5 or reads containing adapter or ploy-N were removed. Paired-end reads were mapped to the cynomolgus reference genome (Macaca_fascicularis_5.0) using HISAT2 (v2.0.5) and quantified using FeatureCounts (v1.5.0-p3).^17, 18^

### Statistical Analysis

Statistical analyses were performed using R (v4.2.2).^19^ Histological data are presented as a boxplot with the median and 25th/75th percentiles. The histological data were analyzed using a linear mixed effects model implemented by “nlme” R package (v3.1-162) followed by Bonferroni correction. Bulk RNA sequencing data were analyzed using a negative binomial mixed effects model with “glmmSeq” R package (v0.5.5).^20^ Dispersion parameter estimates and normalization factor calculation using the trimmed mean of M-value (TMM) method were performed by “edgeR” Bioconductor package.^21^ Gene ontology and Kyoto Encyclopedia of Genes and Genomes (KEGG) pathway enrichment analyses were performed using Enrichr.^22^ Considering correlations among four regions within individual animals, random effects were incorporated in both histological and bulk RNA sequencing data analyses. Correlations between elastic lamellar distance and NOTCH3 area were determined by Spearman method since the data did not pass a normality test. P<0.05 or Bonferroni/false discovery rate (FDR) adjusted P<0.05 was considered statistically significant.

### Data Availability

All numerical data are available in a **Supplemental Excel File**. Bulk RNA sequencing data (raw FASTQ and aligned data) are publicly available at the gene expression omnibus repository (GSE227434).

## RESULTS

### Homogeneous distribution of endothelial cells in the aortic intima and vasa vasorum

We first performed immunostaining of CD31 to evaluate the regional heterogeneity of endothelial cells (ECs, **Supplemental Figure 3A, B**). ECs uniformly formed a single layer in the intima in all aortic regions. In addition to the intima, ECs were also observed in vasa vasorum in the adventitia across the regions. Vasa vasorum were detected in the adventitia, but not in the media. The presence of vasa vasorum into the media was not observed in any regions. These results indicate the uniformity of EC distribution in the aortic wall.

### Unique structural characteristics of elastic fibers in the infrarenal abdominal aorta

We next assessed the aortic media with a focus on SMCs and elastic fibers, by αSMA immunostaining and Verhoeff’s iron hematoxylin staining (**Figure 1A, B**). The aortic size demonstrated by the measurement of the internal perimeters of tissue sections was largest in the ascending aorta (**Supplemental Figure 4A**). The aortic media was also thickest in the ascending aorta (**Figure 1D, Supplemental Figure 4B**), although SMCs were uniformly distributed across the media throughout the aorta (**Figure 1A**). Consistent with the gradient of medial thickness, the ascending aorta had more elastic fibers than those of other regions (**Figure 1E**). The number of elastic fibers was approximately 60 and 10 in the ascending and infrarenal abdominal aortas, respectively. Elastic fibers in the ascending, descending, and suprarenal aortas were distributed densely, whereas those in the infrarenal region were more separated than other regions (**Figure 1F**). Elastic fiber thickness was thinner in the infrarenal region than the ascending and suprarenal regions (**Figure 1G**). In addition, the interlamellar distance of elastic fibers was higher in the infrarenal region compared to other regions (**Figure 1H**), indicating the dispersion of elastic fibers.

**Figure 1.**
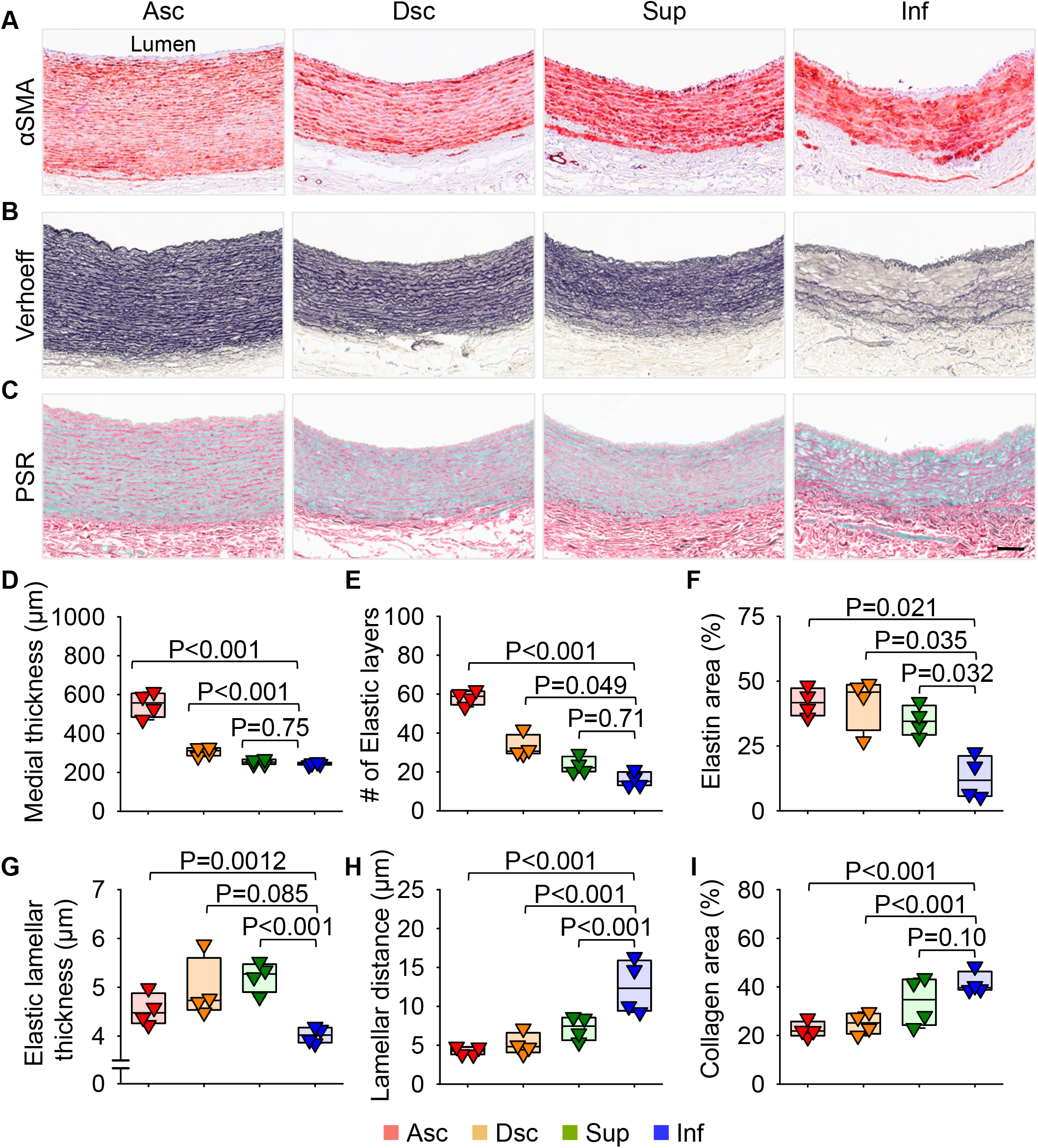
Structural differences of aortic media in young cynomolgus monkeys. Representative images of **(A)** immunostaining for α smooth muscle actin (αSMA), **(B)** Verhoeff’s iron hematoxylin, and **(C)** picrosirius red (PSR) staining. Quantification of **(D)** medial thickness, **(E)** the number of elastic fibers, **(F)** percent elastic area, **(G)** elastic lamellar thickness, **(H)** interlamellar distance, and **(I)** percent collagen area in the media in each region. n = 4 per aortic region. Asc indicates ascending; Dsc, descending; Sup, supra-renal; Inf, infra-renal aorta. Scale bars = 100 μm. P values were calculated using a linear mixed effects model with inverse variance weights followed by Bonferroni correction.

Picrosirius red (PSR) staining was performed to evaluate collagen fibers (**Figure 1C**). Collagen fiber deposition in both the media and adventitia had a gradient that increased from the ascending to infrarenal regions (**Figure 1I**). Significant collagen deposition was also evidenced as indicated by Movat’s staining (**Supplemental Figure 5A**). It is of note that, in the media of the infrarenal aorta, collagen deposition predominated in the area having disarrayed elastic fibers (**Figure 1B, C, Supplemental Figure 5B**). In addition, proteoglycan deposition was observed in the intima adjacent to collagen deposition and disarrayed elastic fibers in the media (**Supplemental Figure 5B**). These results suggest that, compared to other aortic regions, the infrarenal aorta of healthy young monkeys exhibits a disorganized ECM structure characterized by deposition of collagen and proteoglycan, in addition to elastic fiber disruption.

### Distinct transcriptome in the infrarenal abdominal aorta

To explore the regional heterogeneity of aortic transcriptomes, we performed bulk RNA sequencing using aortic mRNA isolated from each region. Principal component analysis revealed that aortic transcriptomes were different in a region-specific manner (**Figure 2A**). In particular, the infrarenal abdominal aorta exhibited a distinct transcriptome compared to other regions (**Figure 2A, B**). There were 132 differentially expressed genes (DEGs, **Figure 2B, Supplemental Excel File**). Many DEGs were associated with regionalization and body axis in a gene ontology analysis (**Supplemental Figure 6A, C**). Reactome pathway enrichment analysis using these DEGs revealed the involvement of collagen-related pathways (**Supplemental Figure 6B, D**).

**Figure 2.**
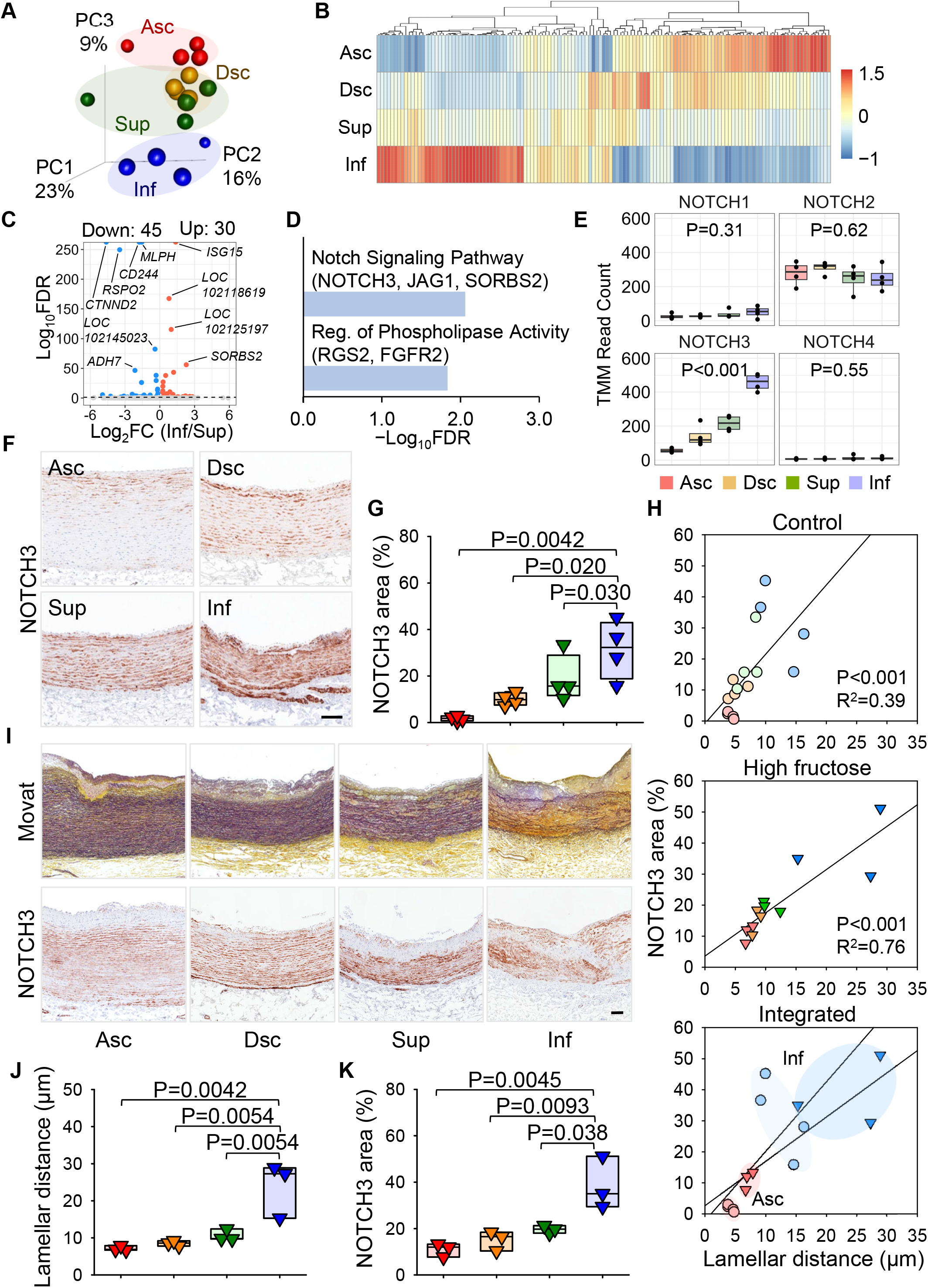
Region-specific transcriptomes and the association of NOTCH3 with elastic fiber dispersion in the infrarenal aorta of cynomolgus monkeys. **(A)** Principal component analysis for unfiltered transcriptomes. **(B)** Heatmap with Z-scored coloring for overall differentially expressed genes (DEGs) identified based on the analysis of deviance. **(C)** Volcano plot for the comparison between infrarenal (Inf) vs suprarenal (Sup) regions and **(D)** gene ontology enrichment analysis for upregulated DEGs. **(E)** Box plots for mRNA read counts of NOTCH family members. **(F)** Immunostaining for NOTCH3 in aortas from female control monkeys (n=4). **(G)** Percent NOTCH3 positive area in the media. **(H)** Linear regression between NOTCH3 and elastin areas in the media of female control monkeys, male adult monkeys provided with high fructose diet and drink, and their integrated data specifically for ascending (Asc) and descending (Dsc) regions. **(I)** Movat’s and immunostaining for NOTCH3 in aortas from male adult monkeys provided with high fructose diet and drink for 3 years (n=3). **(J)** Interlamellar distance of elastic fibers and **(K)** Percent NOTCH3 positive area in the media of male adult monkeys provided with high fructose diet and drink. Scale bar = 100 µm.

Consistent with histological findings that ECs and SMCs were distributed uniformly in the aorta across the regions, mRNA abundances of EC (*PECAM1, VWF, CDH5, TEK*) and SMC (*ACTA2, MYH11, CNN1, TAGLN*) marker genes did not differ among aortic regions (**Supplemental Figure 7A**). Cell proliferation markers, such as *MKI67, MYC, KLF4*, and *KLF5*, were also not different among regions (**Supplemental Figure 7B**). By immunostaining, Ki67 positive cells were rarely detected in any regions (**Supplemental Figure 8**). Although *ELN* and *FBN1* mRNA abundance were not different among regions **(Supplemental Figure 7C)**, some collagen genes showed region-specific differences (**Supplemental Figure 7D**). In the ascending aorta, collagen type 4 A3 and 21 A4 (*COL4A3, COL21A1*) were more abundant than in the infrarenal aorta, whereas collagen type 1 A1 and 6 A3 (*COL1A1, COL6A3*) were more abundant in the infrarenal aorta. Western blot analysis and immunostaining validated the high abundance of collagen type 1 in the infrarenal aorta (**Supplemental Figure 9A, B**).

There is compelling evidence that the renin-angiotensin system and transforming growth factor (TGF-β) signaling affect vascular fibrosis.^23-25^ Therefore, we assessed molecules related to these pathways (**Supplemental Figure 7E, F**). Major molecules in the renin-angiotensin system, *AGT* (angiotensinogen), *ACE* (angiotensin-converting enzyme), and *AGTR1-2* (angiotensin II type 1 or 2 receptor) were not different. mRNA abundance of TGF-β ligands (*TGFB1-3*) and receptors (*TGFBR1-2, LRP1*) were also not different among regions. However, Western blot analysis demonstrated that SMAD2 phosphorylation was significantly lower in the infrarenal aorta compared to other regions, although ERK phosphorylation did not differ among regions (**Supplemental Figure 10A, B**). Given the important role of TGF-β signaling in maintaining the integrity of ECM,^26^ the low TGF-β activity in the infrarenal aorta may contribute to the dispersion and disruption of elastic fibers. We also evaluated vascular inflammation and proteolysis. Some CD68 positive cells were observed in the intima and adventitia, but it was not different across regions (**Supplemental Figure 11**). MMP9 was barely detected in any regions, whereas the cleaved/latent MMP2 ratio in the infrarenal region was higher than that in the ascending aorta (**Supplemental Figure 12**).

To further investigate the molecular basis of regional heterogeneity, transcriptomes were compared between adjacent regions. There were 59 and 27 DEGs in comparisons between ascending vs descending and descending vs suprarenal regions, respectively (**Supplemental Figure 13A, B**), but there was no significant annotation for these genes in gene ontology analyses. Seventy five genes (Up: 30, Down: 45 genes) were different between suprarenal and infrarenal regions (**Figure 2C**). While no term was enriched for downregulated DEGs, Notch signaling pathway was the most significant term in upregulated DEGs (**Figure 2D**). Among Notch family members, only NOTCH3 differed among regions (**Figure 2E**). *NOTCH3* mRNA was most upregulated in the infrarenal region compared to other regions. Immunostaining also demonstrated upregulation of NOTCH3 in the media of the infrarenal aorta (**Figure 2F, G**). Of note, NOTCH3 protein abundance in the media was positively correlated with the interlamellar distance of elastic fibers (**Figure 2H**), consistent with a contribution of NOTCH3 to the elastic fiber dispersion in the infrarenal aorta.

### Augmentation of elastic fiber dispersion and medial NOTCH3 in the infrarenal abdominal aorta of adult monkeys fed high fructose

Region-specific differences of ECM structure and NOTCH3 was observed in young healthy female monkeys. Thus, we next examined whether the regional heterogeneity of these features was augmented in adult male monkeys with pathological stimulation. Atherosclerosis and aortic aneurysm have a regional heterogeneity with the infrarenal region being especially prone,^27, 28^ and “fast food diet” consumption is associated with atherosclerosis.^29^ Therefore, male cynomolgus monkeys at 5 years of age were provided a high fructose diet and drink to mimic a “fast food and soft drink” diet and were terminated 36 months later.

Intimal thickening was observed by Verhoeff iron hematoxylin staining, indicating atherosclerosis formation (**Supplemental Figure 14A**). The structure of aortic ECM, especially elastic fibers, was significantly disrupted in these monkeys. The interlamellar distance of elastic fibers in the infrarenal region was 15 to 29 µm which was higher than that of young female monkeys (**Figure 1H, 2I, J, Supplemental Figure 14A**). Immunostaining of αSMA demonstrated the loss of SMCs in the media of the suprarenal and infrarenal regions (**Supplemental Figure 14B**). Meanwhile, CD68 positive area was observed in the intima uniformly across the regions (**Supplemental Figure 14C**). Similar to young female monkeys, NOTCH3 immunostained area was greatest in the infrarenal aorta (**Figure 2I, K**). Linear regression plots revealed that the association between NOTCH3 and elastic fiber dispersion was enhanced in adult male monkeys (**Figure 2H**).

## DISCUSSION

The present study used histological and transcriptomic approaches to profile aortic cells and ECM in four aortic regions of young cynomolgus monkeys. The infrarenal abdominal aorta displayed disarrayed elastic fibers with collagen and proteoglycan deposition. Transcriptomic analysis identified significantly increased aortic *NOTCH3* that was positively correlated with elastic fiber dispersion. Those features were enhanced in adult male monkeys fed a “fast food” diet. This study highlights the regional heterogeneity of ECM structure and transcriptomes in aortas of non-human primates.

ECM plays a crucial role in maintaining the structural integrity and functions of the aorta. Elastic and collagen fibers are key components of ECM and these fibers collaborate to provide elasticity and strength to the vessel wall.^2, 30, 31^ Thus, the disruption of these fibers could contribute to aortic diseases, such as aneurysms and dissections. This premise is enhanced by fragmentation of elastic fibers and collagen deposition being prominent features of these diseases.^32-34^ In the present study, we found fragmented elastic fibers in the media and collagen deposition in the media and adventitia of the infrarenal abdominal aortas, but not other regions. Thus, these results suggest a structural frailty of the infrarenal aorta. Previous studies reported similar results that elastic fibers were disarrayed and collagen was deposited in the infrarenal aorta of young monkeys.^35, 36^ Interestingly, these histological features were more pronounced in adult monkeys. In the present study, the augmentation of ECM disruption was also observed in aortas of adult monkeys complicated with atherosclerotic lesions. Therefore, the disruption of elastic fibers may be involved in the process of some types of aortic diseases.

Multiple studies have shown that structural components of the aortic wall vary by region. Wolinsky and Glagov compared medial structure between thoracic and abdominal aortic regions in multiple species, including mouse, rat, rabbit, monkey, and human.^37^ Medial thickness and elastic fibers were less abundant in the abdominal region than in the thoracic region across all species. Another study reported consistent results that medial thickness decreases towards the distal region in porcine aortas.^38^ A recent study investigated the difference in elastin and collagen composition in human aortas.^7^ Elastic fibers became less and collagen fibers became more abundant in distal regions. Consistent with these data, the present study identified thinner medial layers, elastin disorder, and higher collagen deposition in the infrarenal aorta. Given the important role of ECM in the integrity of the aortic wall,^26^ the region-specific ECM disruption may contribute to the regional specificity of aortic diseases.

There is evidence that TGF-β signaling contributes to ECM synthesis during aortic development.^39^ A number of studies have reported upregulation of aortic TGF-β signaling associated with elastic fiber fragmentation in many types of aortic aneurysms in humans and mice.^40-43^ Several studies have shown that TGF-β neutralization on aortic aneurysm formation exacerbates aortic aneurysm formation in mice.^44-46^ These support the notion that TGF-β plays a pivotal role in the maintenance of aortic wall and elastic fiber integrity. In the present study, aortic SMAD2 phosphorylation and elastic fiber density were lower in the infrarenal aorta compared to other regions. It is possible that the lower TGF-β signaling is associated with elastic fiber dispersion in the infrarenal aorta.

To determine mechanisms that may contribute to elastic fiber dispersion, the present study examined aortic proteolysis and inflammation. While macrophage accumulation was not different among regions, aortic MMP2 activity was higher in the infrarenal region compared to the ascending region. Importantly, active MMP2 was not different between descending and infrarenal regions. Since elastic fibers in the descending aorta were not disrupted, MMP2 activity may not be a key driving molecule for elastic fiber disruption in the infrarenal aorta. In the bulk RNA sequencing, we found an increase of NOTCH3 in the infrarenal aorta, which was correlated with elastic fiber dispersion. A recent study has reported that genetic deletion of elastin increases aortic NOTCH3 in mice.^47^ Thus, in the present study, increased NOTCH3 may be considered as a consequence of elastic fiber disruption. However, another study reported that pharmacological inhibition of Notch signaling by a γ-secretase inhibitor attenuated elastic fragmentation in Marfan syndrome model mice.^48^ Therefore, further study is needed to unlock the mechanism how NOTCH3 exerts a role in elastic fiber disruption.

Multiple studies have shown that, in addition to elastic fibers, SMCs also play a pivotal role in aortic integrity.^49^ For instance, SMC proliferation and neointimal formation are involved in aortic aneurysms and atherosclerosis formation.^50, 51^ Considering the regional heterogeneity of these diseases, SMCs may have different biological behaviors among regions, which could contribute to the susceptibility of the infrarenal aorta to these diseases. The present study performed bulk RNA sequencing to explore the molecular basis of elastic fiber differences among the four aortic regions. Given the cellular heterogeneity and intercellular interaction, single-cell RNA sequencing would provide more specific insights into understanding molecular mechanisms, including cellular behaviors and cell-cell interaction of SMCs in development and maintenance of elastic fibers.

Selected hereditary connective tissue diseases, such as Marfan syndrome, Loeys-Dietz syndrome, and Ehlers-Danlos syndrome, often develop aortic aneurysms and dissections in the prepubescent phase.^52-54^ Thus, it is important to understand the aortic physiology in the prepubescent phase. In addition, prepubescents have less hormonal variation compared to adults, which enables the determination of aortic functions in a more standardized hormonal environment. In contrast, age and male sex are risk factors for aortic diseases, including abdominal aortic aneurysms. To investigate mechanisms underlying these risk factors, optimal groups, such as age- or sex-matched controls, are required. In the present study, the dispersion of elastic fibers in the infrarenal aorta was assessed in not only young but also adult monkeys. However, these monkeys were not sex-matched. Thus, future studies using age- and sex-matched samples are needed to investigate how the integrity of elastic fibers is maintained in the aorta.

Vasa vasorum form a microvascular network that supplies oxygen and nutrients to the vascular wall.^55-57^ They are also linked to development of vascular diseases, including aortic aneurysms.^58, 59^ Previous studies investigating vasa vasorum formation in multiple species reported that vasa vasorum were observed in the media from the ascending to the suprarenal abdominal aorta, but not in the infrarenal abdominal aorta, in humans.^13, 37, 60^ Based on the number of elastic fibers in each aortic region, this study concluded that medial vasa vasorum in mammals was not present in the aortic wall having less than 29 elastic layers. It has been proposed that the absence of medial vasa vasorum in the infrarenal aorta contributes to the regional heterogeneity of aortic aneurysms.^58, 59^ In our study, vasa vasorum was identified in the adventitia throughout the aorta. In contrast, medial vasa vasorum was not observed in any aortic regions, including the ascending aorta that had more than 29 elastic layers. Therefore, while this may be a variance form previous studies, it should be noted that the present study examined vasa vasorum in young monkeys (3 years old = equivalent to 9 years old in humans) corresponding to the pubescent age in humans,^61^ whereas the previous studies used aortas at 20-50 years of age.^60^ Therefore, vasa vasorum has the potential to change its distribution during aging. Further studies are needed to investigate the determinants of the distribution of vasa vasorum in the aorta.

Studies using non-human primates are time- and cost-consuming compared to mouse studies. In addition, experimental approaches to generate some aortic diseases, such as aneurysms and dissections, have not been developed in monkeys. Thus, a technical advance is needed to enable research approaches in studies using non-human primates for aortic aneurysms. However, non-human primates have clinical relevance because of their biological and anatomical similarities to humans. Thus, the present study provides translational insights into further understanding aortic biology. Considering the limited availability of aortas from healthy humans, our study using aortas from healthy non-human primates leads to a greater understanding of aortic functions under physiological conditions.

In conclusion, our study demonstrated the regional heterogeneity of ECM structure and transcriptomic profiles in aortas from cynomolgus monkeys. The infrarenal abdominal aorta displayed elastic fiber dispersion along with an increase of medial NOTCH3 and collagen fibers and proteoglycan deposition.

## NON-STANDARD ABBREVIATIONS

Asc: ascending thoracic aorta
Dsc: descending thoracic aorta
Sup: suprarenal abdominal aorta
Inf: infrarenal abdominal aorta
EC: endothelial cell
SMC: smooth muscle cell
ECM: extracellular matrix
TGF-β: transforming growth factor β

## Sources of Funding

This project was supported by the National Heart, Lung, and Blood Institute of the National Institutes of Health (R01HL139748 and R35HL155649), the American Heart Association SFRN in Vascular Disease (18SFRN33900001 and 33960163), MERIT Award (23MERIT1036341), and the Novo Nordisk Foundation (NNF18OC0033438). The content in this project is solely the responsibility of the authors and does not necessarily represent the official views of the National Institutes of Health.

## Acknowledgments

Histological images were captured using Zeiss Axioscan 7 in the Light Microscopy Core at the University of Kentucky. The graphic abstract was generated using BioRender.

## Disclosures

None

## Notes

### Competing Interest Statement

The authors have declared no competing interest.

### Summary of Updates

The article has been revised to include new data.

