## Supplemental Materials for "Association of NOTCH3 with Elastic Fiber Dispersion in the Infrarenal Abdominal Aorta of Cynomolgus Monkeys"

**Running title:** Elastic Dispersion and NOTCH3 in Monkey Aorta

**Corresponding authors:**
Hong S. Lu
Saha Cardiovascular Research Center, University of Kentucky
741 South Limestone Street
BBSRB, B249
Lexington, KY, 40536, USA.


Hisashi Sawada
Saha Cardiovascular Research Center, University of Kentucky
741 South Limestone Street
BBSRB, B251
Lexington, KY, 40536, USA.


### Major Resources Table

#### Animals (in vivo studies)

| Species | Vendor or Source | Background Strain | Sex | Persistent ID / URL |
|---|---|---|---|---|
| Mauritian Cynomolgus Monkey | DSP | | M and F | N/A |

#### Antibodies

| Target Antigen | Vendor or Source | Catalog # | Working Concentration | Lot # | URL |
|---|---|---|---|---|---|
| αSMA | abcam | ab5694 | 2 µg/mL | GR3263275-11 | https://www.abcam.com/alpha-smooth-muscle-actin-antibody-ab5694.html |
| CD31 | abcam | ab28364 | 3 µg/mL | GR94677-2 | https://www.abcam.com/cd31-antibody-ab28364.html |
| COL1A1 | abcam | ab138492 | 3 µg/mL | GR3370247-5 | https://www.abcam.com/collagen-i-antibody-epr7785-ab138492.html |
| Ki67 | abcam | ab16667 | 0.03 µg/mL | GR341668-9 | https://www.abcam.com/products/primary-antibodies/ki67-antibody-sp6-ab16667.html |
| NOTCH3 | abcam | ab23426 | 1 µg/mL | 1012556-1 | https://www.abcam.com/products/primary-antibodies/notch3-antibody-ab23426.html |
| CD68 | Bio Legend | 916104 | 0.5 µg/mL | B228266 | https://www.biolegend.com/en-us/products/purified-anti-cd68-antibody-13199 |
| pSMAD2 | CST | 3108S | 0.1 µg/mL | 8 | https://www.cellsignal.com/products/primary-antibodies/phospho-smad2-ser465-467-138d4-rabbit-mab/3108 |
| SMAD2 | CST | 3122S | 0.1 µg/mL | 8 | https://www.cellsignal.com/products/primary-antibodies/smad2-86f7-rabbit-mab/3122 |
| pERK | CST | 9101S | 0.1 µg/mL | 29 | https://www.cellsignal.com/products/primary-antibodies/phospho-p44-42-mapk-erk1-2-thr202-tyr204-antibody/9101 |
| ERK | CST | 9102S | 0.1 µg/mL | 27 | https://www.cellsignal.com/products/primary-antibodies/p44- |

| | | | | | |
|---|---|---|---|---|---|
| | | | | | 42-mapk-erk1-2-antibody/9102 |
| β-actin | Sigma-Aldrich | A5441 | 0.3 μg/mL | 079M4799V | https://www.sigmaaldrich.com/US/en/product/sigma/a5441 |
| Goat anti-rabbit | Vector | PI-1000 | 1.0 μg/mL | ZG1009 | https://vectorlabs.com/products/antibodies/peroxidase-goat-anti-rabbit-igg |
| Goat anti-mouse | Sigma-Aldrich | A2554 | 0.3 μg/mL | 175122 | https://www.sigmaaldrich.com/US/en/product/sigma/a2554 |

**DNA/cDNA Clones**

| Clone Name | Sequence | Source / Repository | Persistent ID / URL |
|---|---|---|---|
| N/A | | | |

**Cultured Cells**

| Name | Vendor or Source | Sex (F, M, or unknown) | Persistent ID / URL |
|---|---|---|---|
| N/A | | | |

**Data & Code Availability**

| Description | Source / Repository | Persistent ID / URL |
|---|---|---|
| GSE227434 | GEO | https://www.ncbi.nlm.nih.gov/geo/query/acc.cgi?acc=GSE227434 |

**Other**

| Description | Source / Repository | Persistent ID / URL |
|---|---|---|
| N/A | | |

**Supplemental Table 1. Semisynthetic diet and drink composition for male cynomolgus monkeys**

**Fast food diet**

| Ingredient | | g/100 g dry weight |
|---|---|---|
| Casein | | 9.0 |
| Whey Protein Isolate 90 | | 5.0 |
| Sucrose | | 15.0 |
| Fructose | | 20.0 |
| Whole Wheat Flour | | 20.0 |
| Lard | | 16.4 |
| Fish Oil | | 0.2 |
| Cholesterol | | 0.2 |
| Cellulose | | 6.3 |
| Vitamin Mix, X75 | | 2.5 |
| Mineral Mix, Hegsted IV | | 5.0 |
| Calcium Carbonate | | 0.4 |
| | **TOTAL** | **100.0** |

| Macronutrient | Calories (%) |
|---|---|
| Fat | 37 |
| Carbohydrate | 49 |
| Protein | 14 |

**High fructose drink**

| Ingredient | g/100 g dry weight |
|---|---|
| Fructose (g/L) | 60.5 |
| Glucose (g/L) | 49.5 |
| Tropical Punch Concentrate (mL/L) | 10 |
| Red Food Coloring (mL/L) | 0.2 |
| **Calories/mL** | 0.4 |

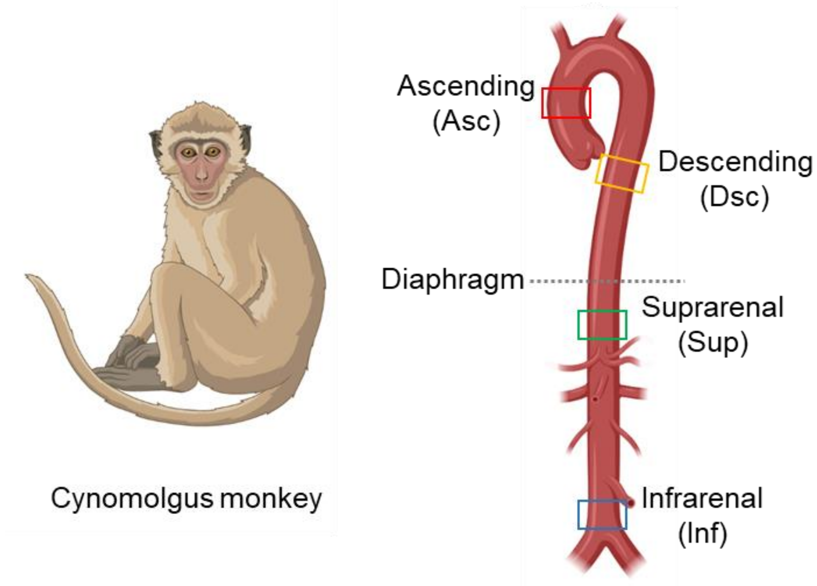


**Supplemental Figure 1. Aortic tissues harvested from cynomolgus monkeys.** Aortic tissues were harvested from the middle ascending aorta (Asc, red box), the middle descending thoracic aorta (Dsc, yellow box), the suprarenal abdominal aorta at the proximal to the celiac artery (Sup, green box), and the infrarenal abdominal aorta proximal to the bifurcation of iliac arteries (Inf, blue box). Figures are created with BioRender.com.

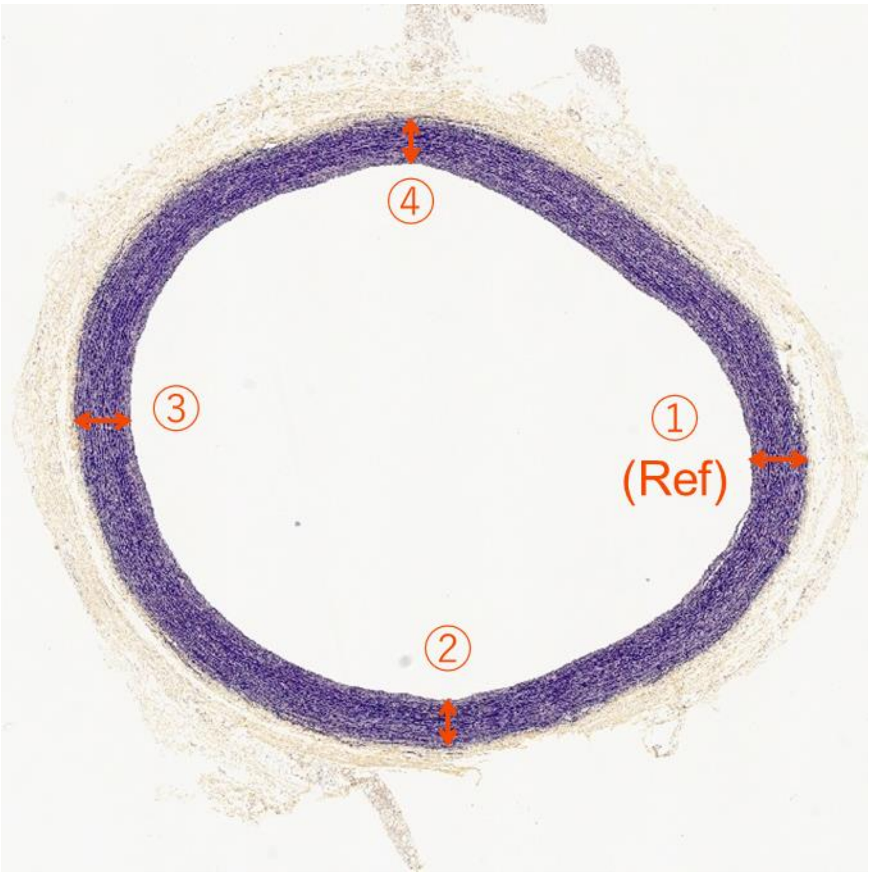


**Supplemental Figure 2. Example image for histological measurements.**
The most thickened media was set as a reference point (Ref). Medial thickness, number of elastic fibers, elastic lamellar thickness, and interlamellar distance of elastic fibers were measured in four orthogonal locations of each section and means were calculated.

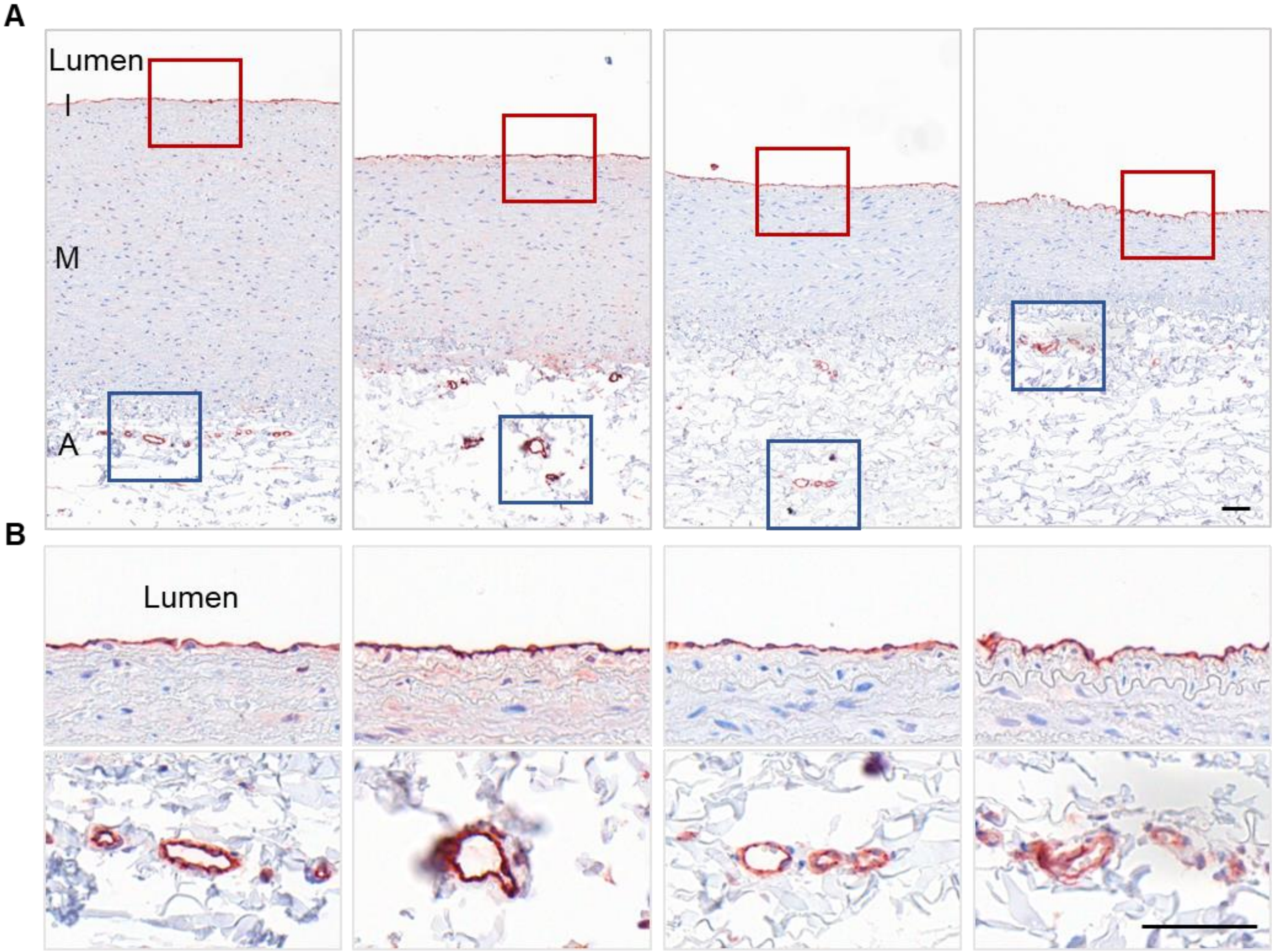


**Supplemental Figure 3. Homogenous distribution of endothelial cells in aortas of cynomolgus monkeys.**
Representative images of immunostaining for CD31 in **(A)** low and **(B)** high magnifications. n=4 per aortic region. Asc indicates ascending; Dsc, descending; Sup, supra-renal; Inf, infra-renal aorta; I, intima; M, media; A, adventitia. Scale bars = 50 µm.

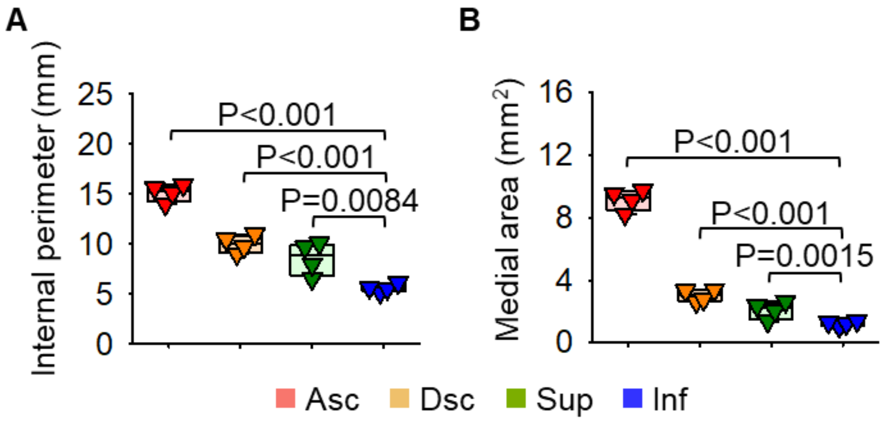


**Supplemental Figure 4.**
**(A)** Internal perimeter length and **(B)** medial area measured in aortic tissue sections from 4 aortic regions (Asc: ascending, Dsc: descending, Sup: suprarenal, Inf: infrarenal) of cynomolgus monkeys. n=4 per aortic region.

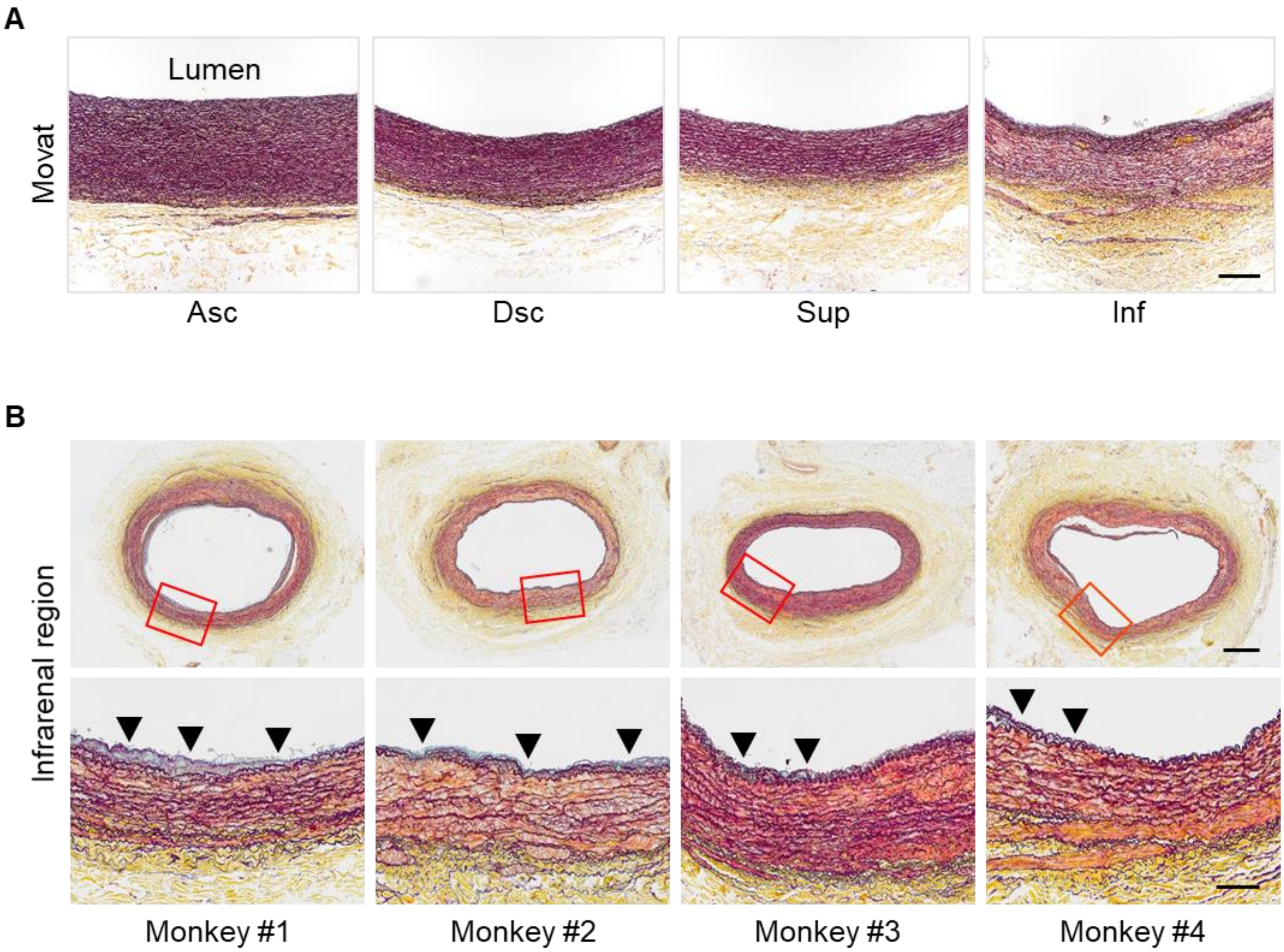


**Supplemental Figure 5. Disruption of the extracellular matrix in the media of the infrarenal aorta from cynomolgus monkeys.** Representative images of Movat's staining from **(A)** 4 aortic regions and **(B)** the infrarenal aorta of 4 monkeys. Asc indicates ascending; Dsc, descending; Sup, supra-renal; Inf, infra-renal aorta. Black triangles indicate proteoglycan deposition. Scale bars = 100 or 400 μm.

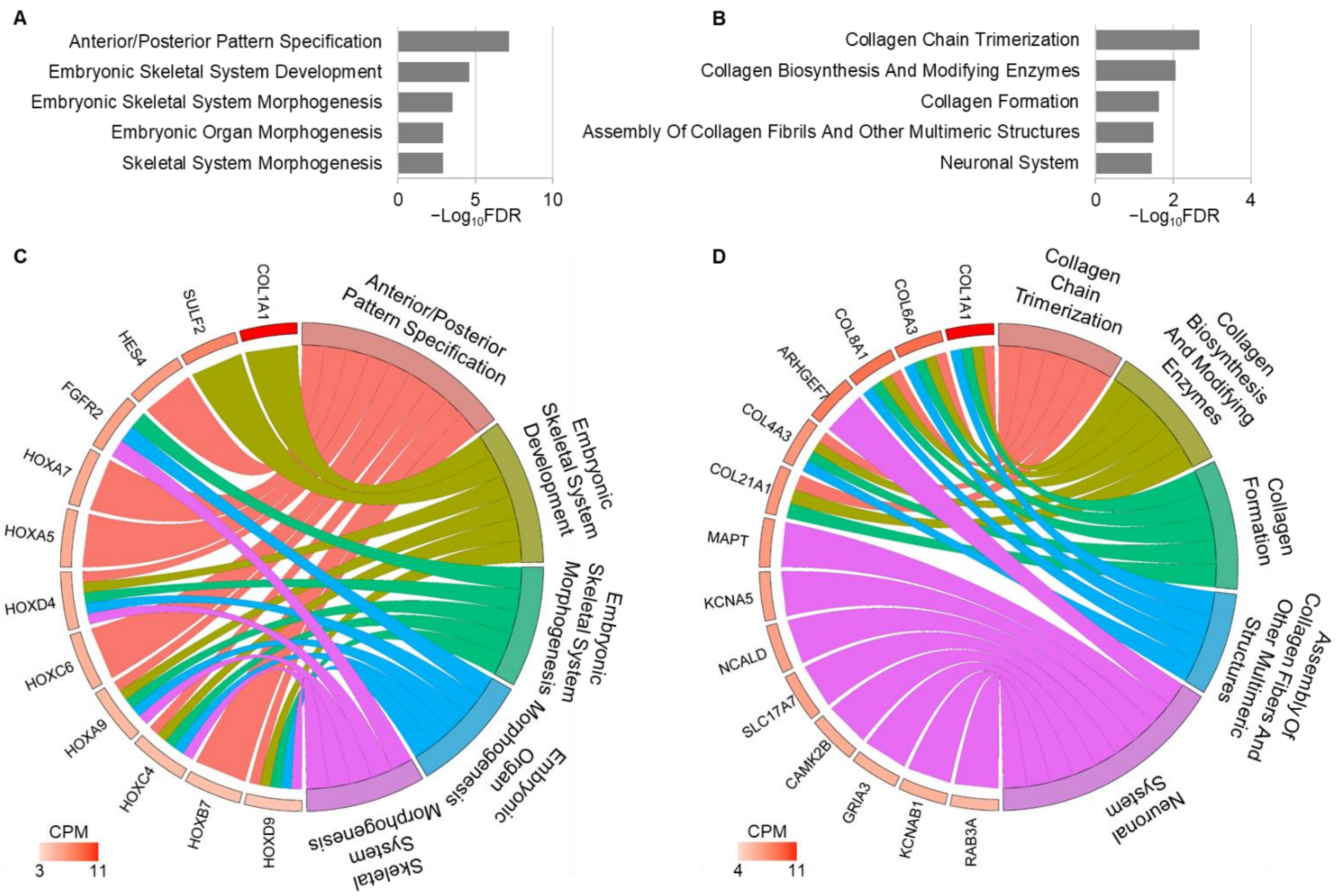


**Supplemental Figure 6. Enrichment analyses for the biological process in gene ontology and the Reactome pathway.**

Top 5 annotations in enrichment analyses for **(A)** the biological process and **(B)** Reactome pathway using differentially expressed genes (DEGs) among regions. Chord graph representing DEGs corresponding to the top 5 terms of **(C)** the gene ontology and **(D)** Reactome pathway. The gene name color code represents the count per million (CPM).

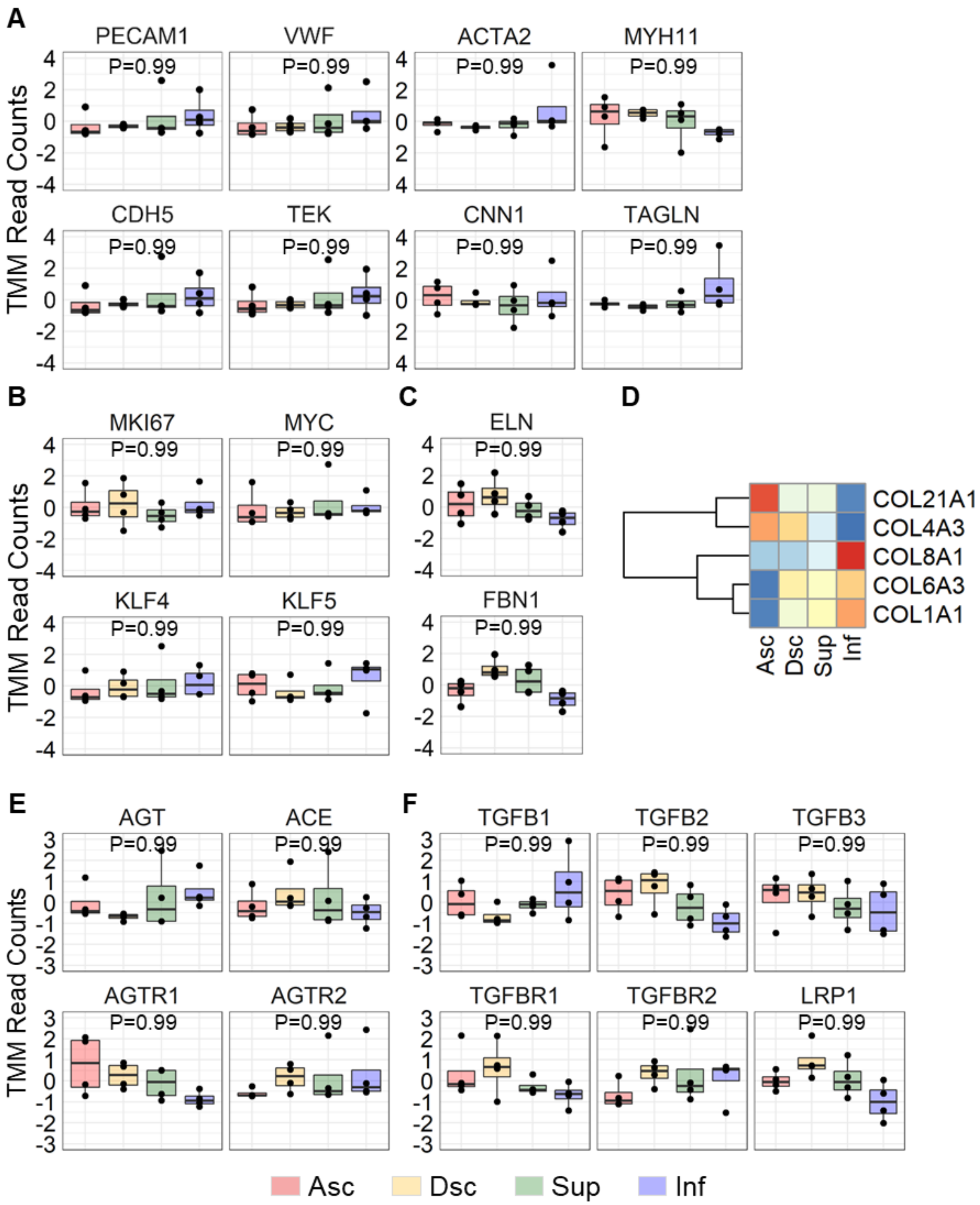


**Supplemental Figure 7. Box plots and a heat map for featured genes.**
Box plots for genes related to **(A)** cell markers for endothelial and smooth muscle cells, **(B)** cellular proliferation, **(C)** elastin (*ELN*) and fibrillin-1 (*FBN1*), **(E)** major components of the renin angiotensin system (RAS), and **(F)** transforming growth factor (TGF-β) ligands and receptors. **(D)** Heat map for differentially expressed collagen genes. n=4 per aortic region. Asc indicates ascending; Dsc, descending; Sup, supra-renal; Inf, infra-renal aorta.

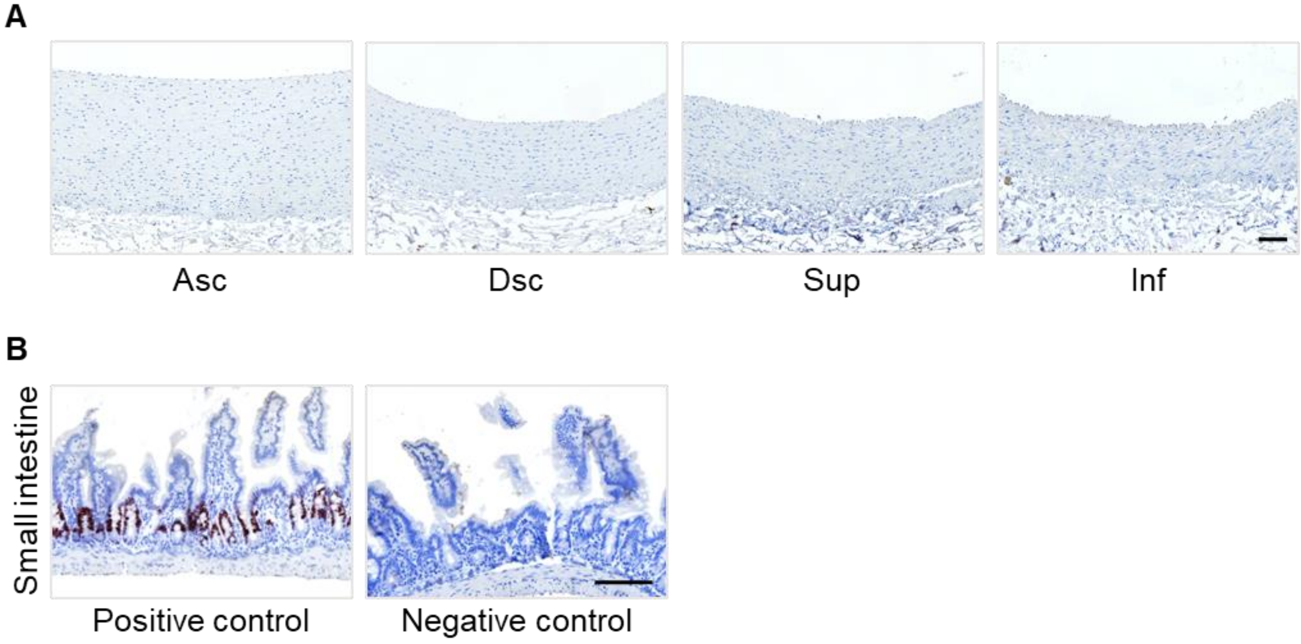


**Supplemental Figure 8. Immunostaining of Ki67 in the aorta of young cynomolgus monkeys.**
Representative images for immunostaining of Ki67 in **(A)** 4 aortic regions from young control monkeys (n=4 per aortic region) and **(B)** small intestines of a wild-type mouse. The mouse intestine was also stained with isotype-match non-immune IgG as a negative control. Asc indicates ascending; Dsc, descending; Sup, suprarenal; Inf, infrarenal aorta.

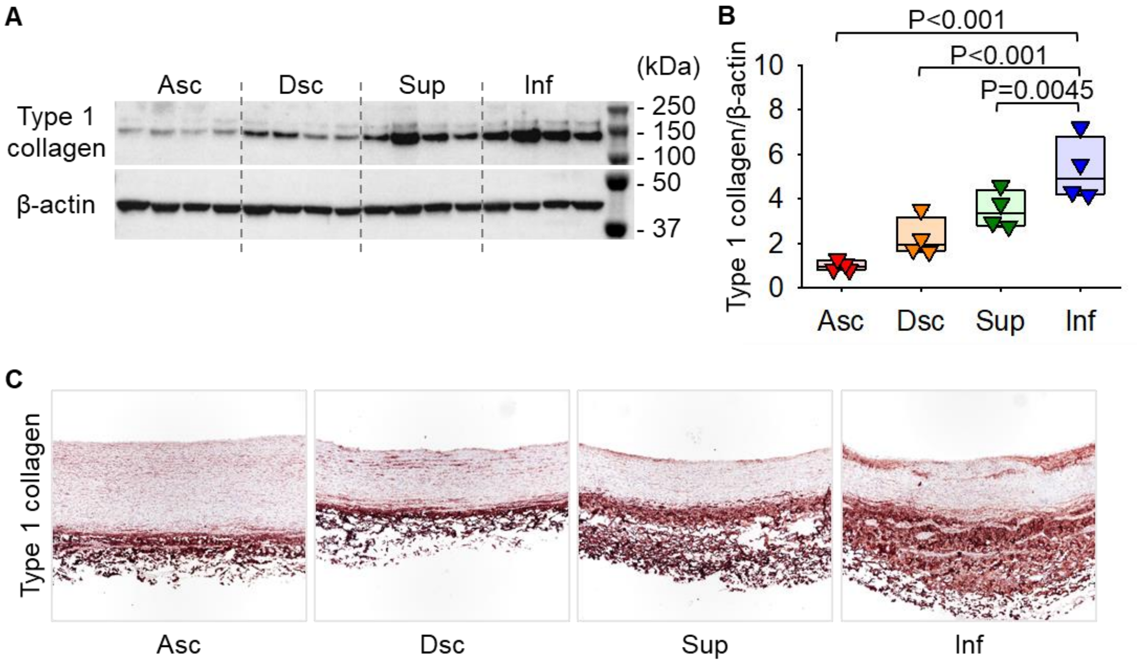


**Supplemental Figure 9. Transaortic gradient of type 1 collagen in cynomolgus monkeys. (A)** Western blot for type 1 collagen and β-actin in aortas of young cynomolgus monkeys and **(B)** its quantification. **(C)** Representative images of immunostaining for type 1 collagen. Asc indicates ascending aorta; Dsc, descending thoracic aorta; Sup, suprarenal aorta; Inf, infrarenal aorta.

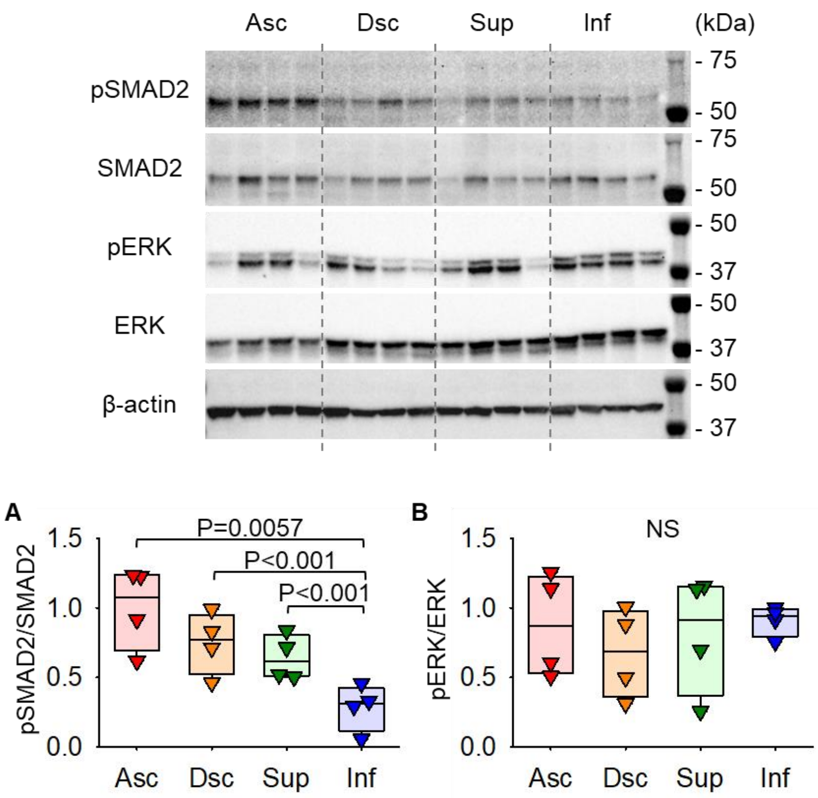


**Supplemental Figure 10. Regional heterogeneity of aortic TGF-β activity in cynomolgus monkeys.**

Western blot analysis for **(A)** SMAD2 and **(B)** ERK phosphorylation in 4 aortic regions. n=4 per aortic region. Asc indicates ascending aorta; Dsc, descending thoracic aorta; Sup, suprarenal aorta; Inf, infrarenal aorta; NS, not significant.

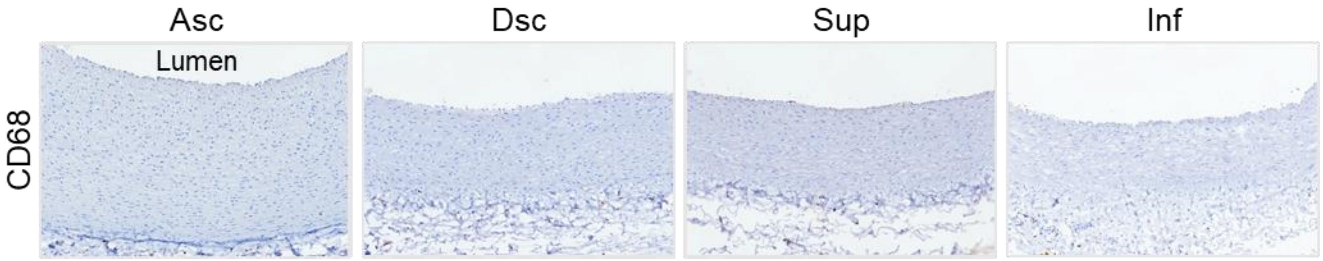


**Supplemental Figure 11. Aortic macrophage accumulation in young cynomolgus monkeys.** Representative images of immunostaining for CD68. Asc indicates ascending aorta; Dsc, descending thoracic aorta; Sup, suprarenal aorta; Inf, infrarenal aorta.

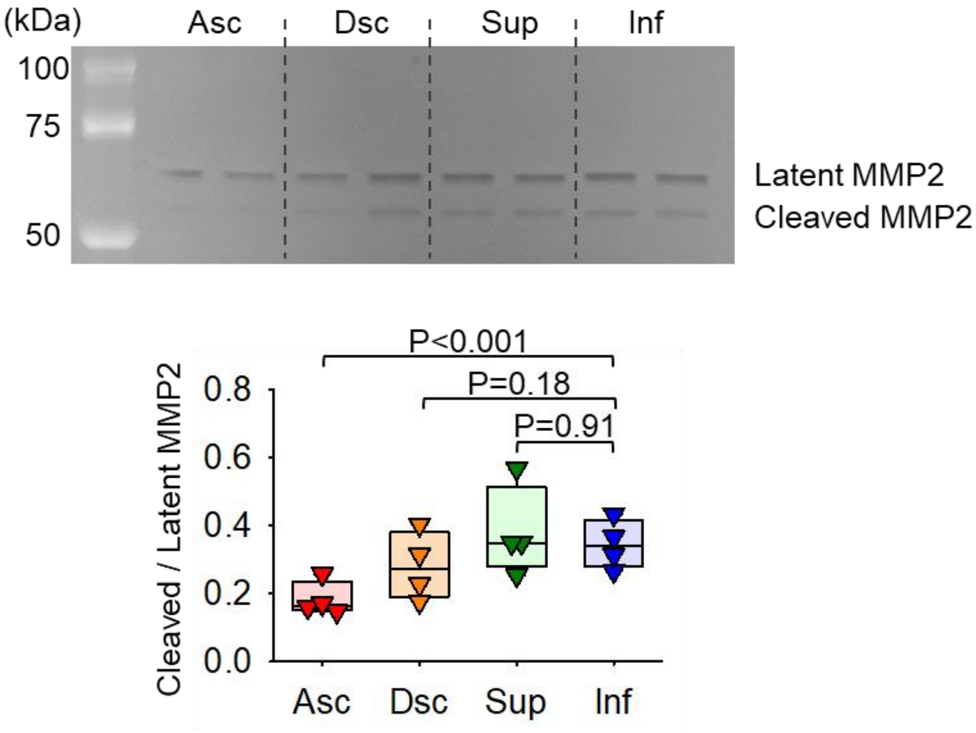


**Supplemental Figure 12. Regional heterogeneity of aortic MMP2 and MMP9 activities in control cynomolgus monkeys.**
A representative gel image of a gelatin zymography and its quantification using 4 aortic regions of young cynomolgus monkeys. n=4 per aortic region. Only latent and cleaved forms of MMP2 were detected. Asc indicates ascending aorta; Dsc, descending thoracic aorta; Sup, suprarenal aorta; Inf, infrarenal aorta.

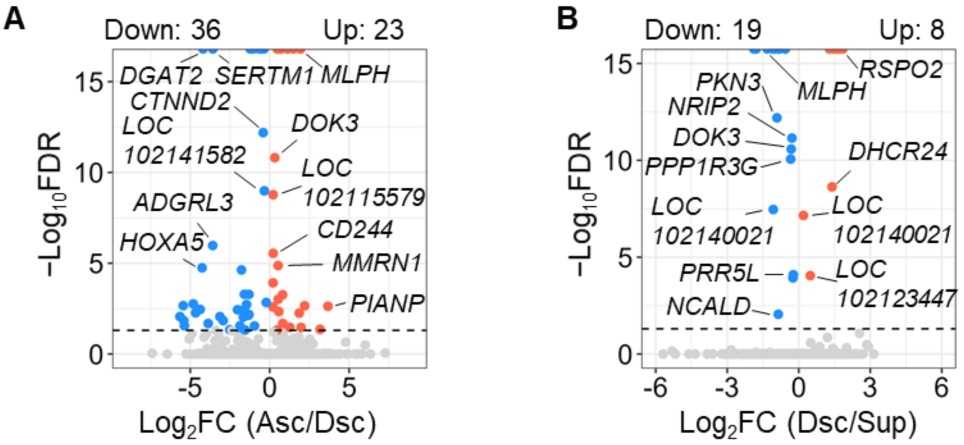


**Supplemental Figure 13. Transcriptomic differences between aortic regions in cynomolgus monkeys.**

Volcano plots for differentially expressed genes in comparing transcriptomes between **(A)** ascending (Asc) and descending (Dsc), and **(B)** Dsc and suprarenal (Sup) aortas. n=4 per aortic region.

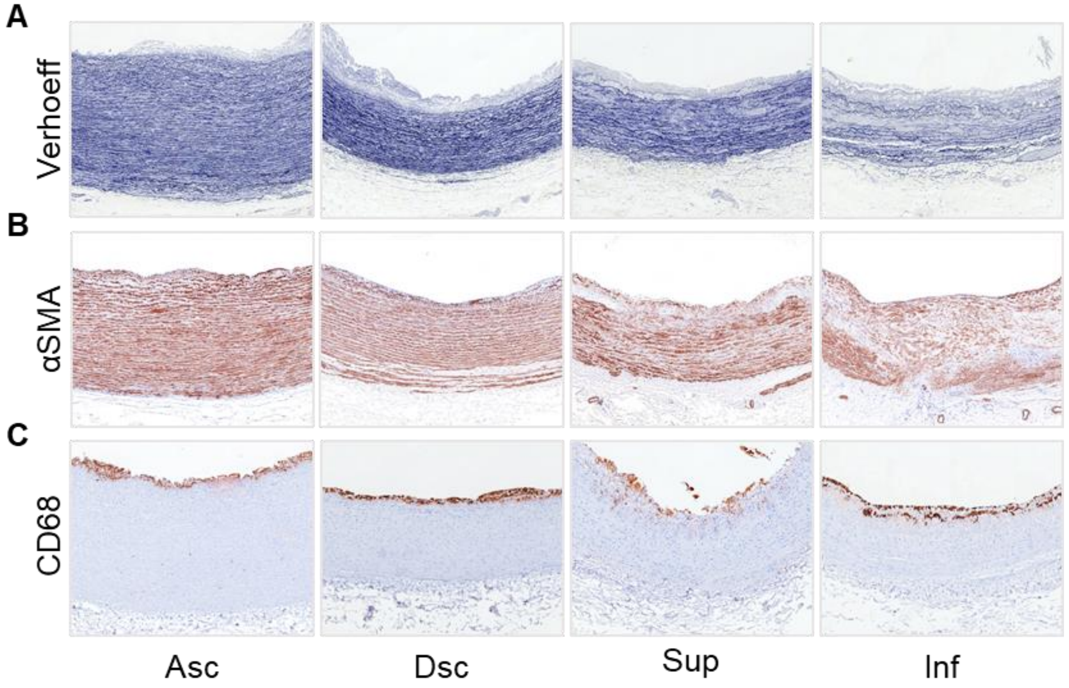


**Supplemental Figure 14. Regional difference of aortic structure in adult cynomolgus monkeys fed with a high fructose diet.**
Representative images of **(A)** Verhoeff iron hematoxylin and immunostaining of **(B)** αSMA and **(C)** CD68 in the 4 aortic regions of cynomolgus monkeys at 8 years of age provided with a high fructose diet and drink for 3 years. n=4 per aortic region. Asc indicates ascending aorta; Dsc, descending thoracic aorta; Sup, suprarenal aorta; Inf, infrarenal aorta
